# Antibody elicited by HIV-1 immunogen vaccination in macaques displaces Env fusion peptide and destroys a neutralizing epitope

**DOI:** 10.1101/2021.03.17.435265

**Authors:** Morgan E. Abernathy, Harry B. Gristick, Jost Vielmetter, Jennifer R. Keeffe, Priyanthi NP Gnanapragasam, Yu E. Lee, Amelia Escolano, Rajeev Gautam, Michael S. Seaman, Malcolm A. Martin, Michel C. Nussenzweig, Pamela J. Bjorkman

## Abstract

HIV-1 vaccine design aims to develop an immunogen that elicits broadly neutralizing antibodies against a desired epitope, while eliminating responses to off-target regions of HIV-1 Env. Here we report isolation and characterization of Ab1245, an off-target antibody against the Env gp120-gp41 interface, from V3-glycan patch immunogen-primed and boosted macaques. A 3.7Å cryo-EM structure of an Ab1245-Env complex reveals one Ab1245 Fab binding asymmetrically to Env trimer at the gp120-gp41 interface using its long CDRH3 to mimic regions of gp41. The mimicry includes positioning of a CDRH3 methionine into the gp41 tryptophan clasp, resulting in displacement of the fusion peptide and fusion peptide-proximal region. Despite fusion peptide displacement, Ab1245 is non-neutralizing even at high concentrations, implying that only two fusion peptides per trimer are required for viral–host membrane fusion. These structural analyses facilitate immunogen design to prevent elicitation of Ab1245-like antibodies that block neutralizing antibodies against the fusion peptide.

## Introduction

Recent efforts in vaccine design for the HIV-1 virus have focused on developing neutralizing adaptive immune responses to the HIV-1 Env glycoprotein via sequential immunization^1–3^. Studies of broadly neutralizing antibodies (bNAbs) isolated from HIV+ human donors have informed immunogen design efforts for various epitopes on the Env trimer, including the V3-glycan patch^4,5^, the fusion peptide (FP)^6,7^, and the CD4-binding site^8,9^. In some cases, on-target antibody responses are accompanied by off-target responses in which antibodies are made against undesired epitopes on the Env trimer including the ‘bottom’ or ‘base’ epitope^10^ and/or a minimally-glycosylated region (glycan ‘hole’)^11^. These antibodies target immunodominant but non-neutralizing epitopes and therefore do not contribute meaningfully to a neutralizing antibody response.

We previously described the design and characterization of RC1, a BG505 SOSIP.664^12^-based engineered immunogen targeting the V3-glycan patch on the gp120 subunit of Env trimer^4^. We showed that RC1 and/or RC1-4fill (modified from RC1 to include additional potential N-linked glycosylation sites; PNGSs) that had been multimerized on virus-like particles (VLPs) elicited antibodies that recognized the V3-glycan patch in wild-type mice, rabbits, and non-human primates (NHPs)^4^. We subsequently boosted a subset of RC1-4fill-primed NHPs, isolated single Env-specific B cells, and derived antibody sequences from which monoclonal antibodies (mAbs) were produced^13^. Here, we describe a single-particle cryo-EM structure of a BG505 Env trimer bound to a monoclonal antibody (Ab1245) isolated from a rhesus macaque after a sequential immunization protocol that included multimerized HIV-1 SOSIP Envs derived from different clades. Ab1245 binds to an epitope overlapping with the FP-targeting bNAb VRC34^6^ at the interface of the Env gp41 and gp120 subunits, but unlike VRC34, Ab1245 displaces the FP and fusion peptide-proximal region (FPPR). In addition, Ab1245 contains a methionine residue that structurally mimics Met530_gp41_, a key residue for the stability of the Env trimer, by engaging the “tryptophan clasp” formed by three gp41 tryptophan residues^14,15^. Despite inducing FP rearrangement and overlap with the neutralizing VRC34 epitope, Ab1245 did not neutralize BG505 or other viral strains, perhaps because of its sub-stoichiometric binding to Env trimer. These previously-unseen features of gp120-gp41 interface antibodies demonstrate that HIV-1 Env can elicit non-neutralizing antibodies that block a neutralizing epitope, inform immunogen design protocols to prevent elicitation of similar antibodies, and provide potential mechanistic insight into HIV-1 Env-mediated fusion of the host and viral membranes.

## Results and Discussion

### Sequential immunization after RC1-4-fill priming elicited Ab1245, a non-V3-targeting antibody

We previously described a V3-glycan patch targeting immunogen, RC1, which was modified from a designed V3 immunogen, 11MUTB^16^, by removing the N-linked glycan attached to gp120 residue N156gp120^4^. Both RC1 and 11MUTB were derived from clade A BG505 SOSIP.664 native-like Env trimers^12^. RC1-4fill and 11MUTB-4fill were modified from RC1 and 11MUTB, respectively, to reduce antibody responses to off-target epitopes^11,17–19^ by inserting PNGSs to add glycans to residues 230_gp120_, 241_gp120_, 289_gp120_, and 344_gp120_^4^. In addition, to enhance avidity effects and limit antibody access to the Env trimer base, we multimerized immunogens on VLPs using the SpyTag-SpyCatcher system^20,21^. Four NHPs primed with RC1-4fill-VLPs^4^ were boosted sequentially with (i) VLPS coupled with 11MUTB-4fill^16^ (clade A), (ii) VLPs coupled with B41 SOSIP (clade B), and (iii) VLPs coupled with a mixture of AMC011 and DU422 SOSIPs (clades B and C) over the course of 9 months. The sequences of the heavy and light chains of Ab1245 were generated by single cell cloning from B cells isolated from one of the boosted NHPs that were captured using BG505 and B41 SOSIPs as baits as described^13^ (Fig. 1a) The heavy and light chains were derived from the macaque V gene segments IGHV4-2*01 and IGLV9-1*01, respectively, and exhibited 14% (heavy chain) and 5% (light chain) amino acid changes due to somatic hypermutation. Of note, the third complementarity region (CDR) of the heavy chain (CDRH3) was longer than typical macaque CDRH3s (24 residues compared with an average of 13-15 residues^22^). Initial binding characterizations of the Ab1245 Fab showed that it bound to BG505 SOSIP and to both RC1 and RC1 glycan KO-GAIA (RC1 lacking the N301_gp120_ and N332_gp120_ glycans surrounding the V3-glycan patch with additional mutations to remove the gp120 GDIR sequence^4^), suggesting that it does not target the V3-glycan patch (Fig. 1b).

**Figure 1.**
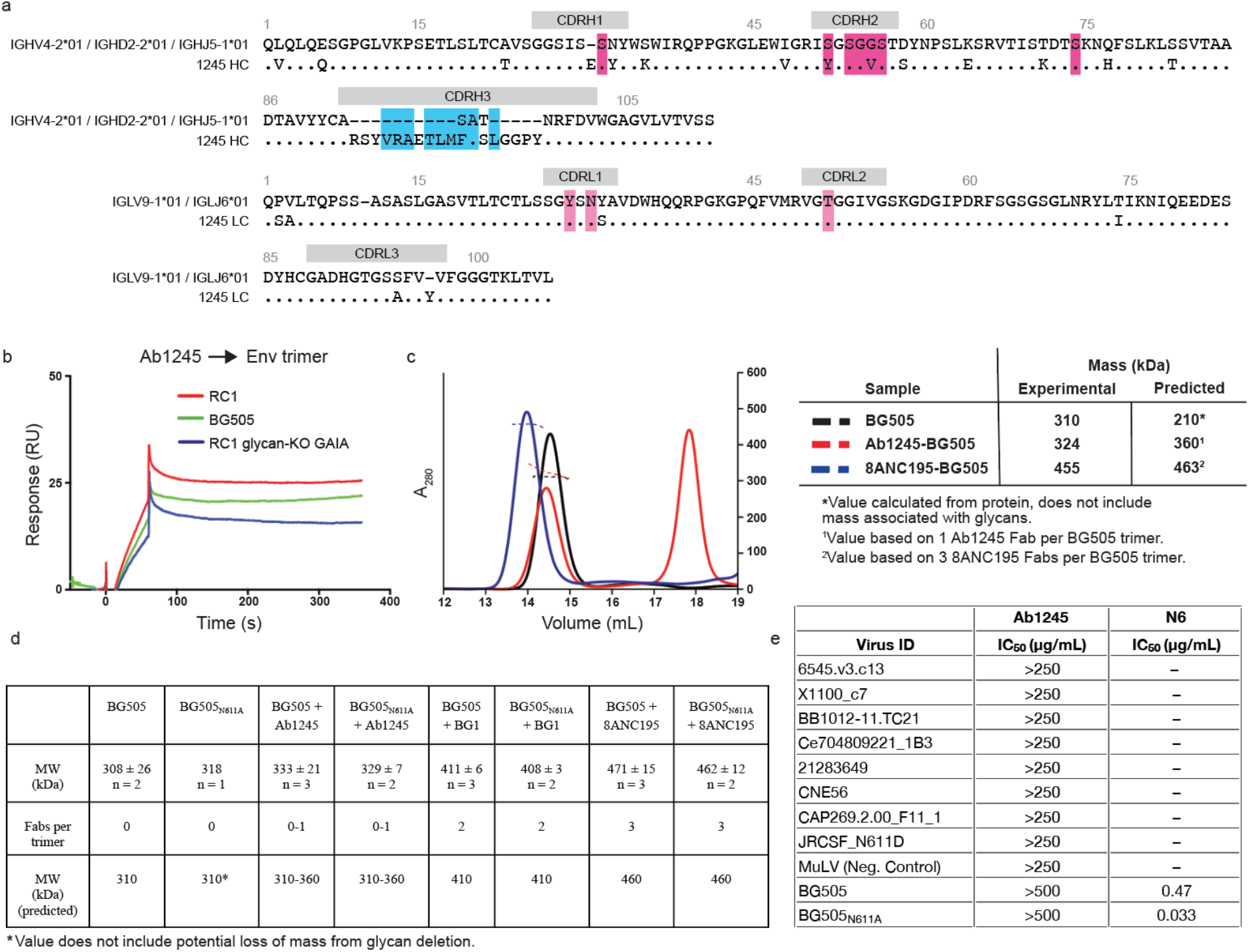
Characterization of Ab1245 elicited in macaques by sequential immunization. **a**, Sequence alignment of Ab1245 heavy and light chains with their germline precursors. Contacts with BG505 Env are indicated by a colored box around the residue (cyan box of CDRH3 contact residues; dark pink and light pink boxes for heavy chain and light chain contact residues, respectively). Residues within CDRs are indicated; residues between CDRs are within framework regions (FWRs). CDRH3 residues derived from VDJ joining are shown as dashes in the top germline sequence, and changes from the germline precursors are denoted a different residue in the mature Ab1245 sequence. Residues are numbered using the Kabat convention. **b**, SPR comparing binding of Ab1245 Fab to RC1, BG505, and RC1 glycan KO-GAIA. **c**, SEC-MALS profiles for BG505 SOSIP.664 Env trimer alone and complexed with a 3-fold molar excess of Ab1245 and 8ANC195 Fabs. Left: absorbance at 280 nm (left y-axis) plotted against elution volume from a Superdex 200 10/300 GL gel filtration column overlaid with the molar mass determined for each peak (right y-axis). Right: Table showing predicted and calculated molecular masses. **d**, Mass photometry results. Derived molecular masses (MW) are listed for Env trimers (either BG505 or BG505_N611A_) incubated without an added Fab or with the indicated Fab (Ab1245, BG1, or 8ANC195) as mean and standard deviation for the indicated number of independent measurements. The Fabs/trimer row shows the expected number of Fabs for each Fab/Env trimer complex. The predicted mass row shows the mass calculated assuming 310 kDa for BG505 trimer (derived by SEC-MALS, panel b) plus 50 kDa per bound Fab. **e**, In vitro neutralization assays using IgGs Ab1245 or N6^62^ (positive control for neutralization) against the indicated viral strains. In addition to those listed, Ab1245 was tested in-house against pseudoviruses from strains CE0217, CNE55, JRCSF, Du422, T250-4, Tro, X1632, 246F3, CH119, CE1176, BJOX002000_03_02, 25710, X2278, CNE8, and 398F1 at a top concentration of 500 μg/mL or 1000 μg/mL along with IgG N6 (positive control at a top concentration of 10 μg/mL). Whereas N6 exhibited expected neutralization potencies^62^, Ab1245 exhibited no neutralization activity.

### Only one Ab1245 Fab binds to each BG505 SOSIP trimer

To determine the binding stoichiometry for the Ab1245 Fab interaction with BG505, we derived the absolute molecular mass of BG505-1245 Fab complexes using size-exclusion chromatography combined with multi-angle light scattering (SEC-MALS). When BG505 was incubated overnight with a 3-fold molar excess of Ab1245 Fab (three Fabs per gp120-gp41 protomer), we observed a heterogeneous mixture corresponding to zero to one Ab1245 Fabs bound per trimer, whereas incubation with 8ANC195 Fab resulted in a homogeneous complex corresponding to three Fabs per trimer (Fig. 1c), as expected from previous stoichiometry measurements and structures^23–25^. To verify that one or more Ab1245 Fabs per trimer did not dissociate during the chromatography procedure required for SEC-MALS, we used mass photometry, a technique that derives approximate masses for individual proteins and complexes in solution^26^, to measure the molecular masses of BG505 alone and complexed with Ab1245 or with control Fabs: 8ANC195 (three Fabs per BG505 trimer)^23–25^ and BG1 (two Fabs per BG505 trimer)^23^. Consistent with the SEC-MALS results, mass photometry experiments suggested zero to one Ab1245 Fabs bound to each wild-type BG505 trimer and to a N611A mutant BG505 trimer (Fig. 1d). We conclude that Ab1245 Fab binds asymmetrically to Env with at most one Fab per trimer.

### Ab1245 binds at the gp120-gp41 interface

To further characterize the Ab1245 epitope on HIV-1 Env, we solved a single-particle cryo-EM structure of Ab1245 Fab bound to a BG505 Env trimer. To form complexes, we incubated a 3-fold molar excess of Ab1245 Fab with BG505, followed by an incubation with a 3-fold excess of 8ANC195^27^ Fab to add mass to the complex and prevent problems associated with preferred orientation bias (Table 1, Supplementary Fig. 1c). This resulted in complex formation with three 8ANC195 Fabs and a maximum of one Ab1245 Fab bound per BG505 trimer (Fig. 2a), consistent with the stoichiometry experiments (Fig. 1c,d). The complex with one Ab1245 Fab per BG505 trimer was solved at 3.7 Å resolution and showed generally well-defined side chain density throughout the complex (Fig. 2b; Supplementary Fig. 1f). An additional 3D class of BG505 trimers was observed with three bound 8ANC195 Fabs and no Ab1245 Fabs (Supplementary Fig. 1e).

**Figure 2.**
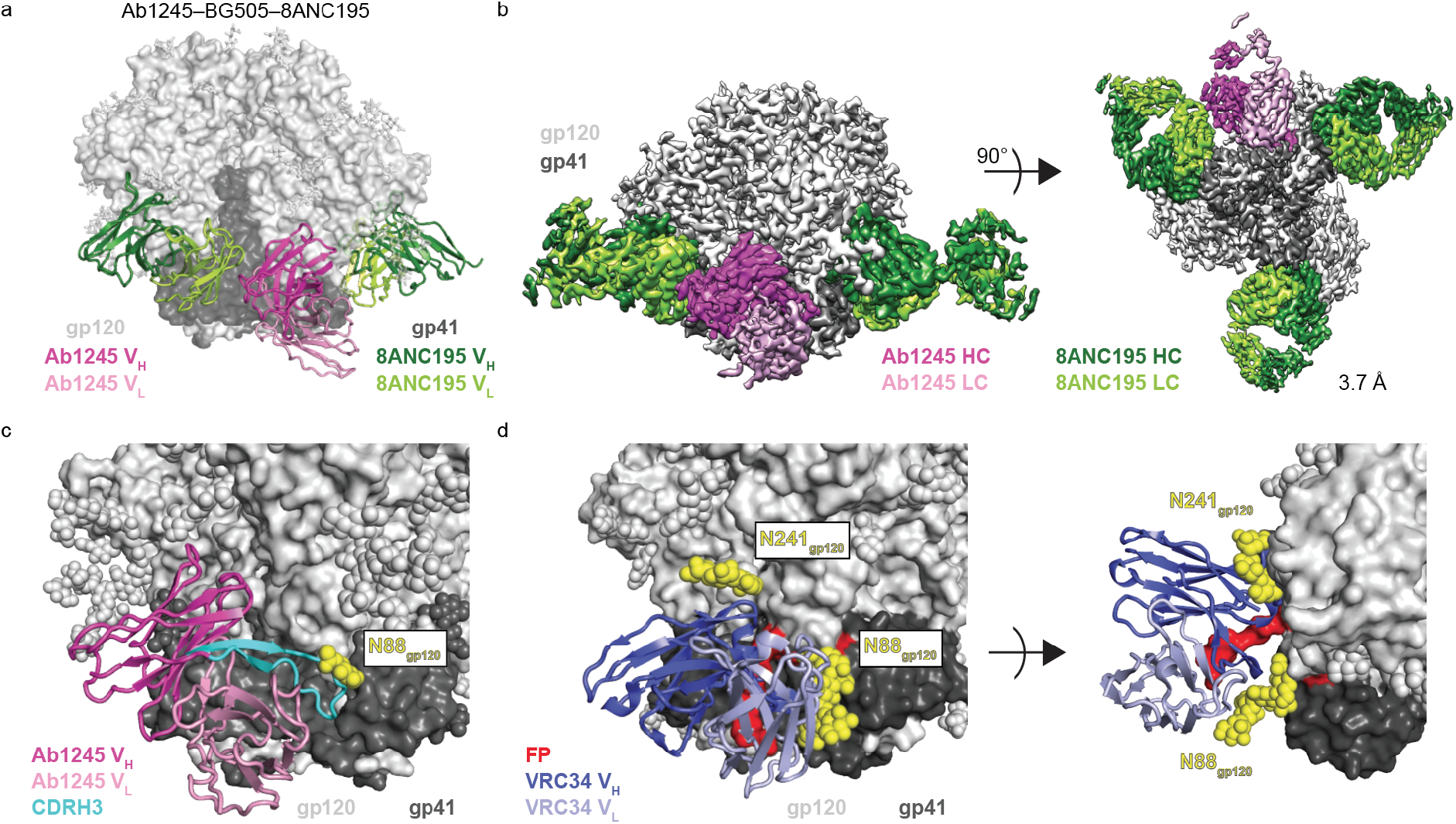
Ab1245 binds the gp120-gp41 interface. **a**, Representation of Ab1245-BG505-8ANC195 structure. Fabs are shown in cartoon, BG505 is shown as surface, and glycans are shown as sticks. Within BG505, gp120 is light gray and gp41 is dark gray. The Ab1245 V_H_-V_L_ is dark pink (heavy chain) and light pink (light chain); 8ANC195 Fabs are dark green (heavy chain) and light green (light chain). **b**, Side view (left) and view looking up from the trimer base (right) of density map for 3.7 Å Ab1245-BG505-8ANC195 complex. Colors as in panel a. **c**, Close-up view of Ab1245 V_H_-V_L_ domains (cartoon) interacting with gp120-gp41 interface with highlighted CDRH3 (cyan). N-linked glycans are light gray (gp120), dark gray (gp41), or yellow (N88_gp120_ and N241_gp120_ glycans) spheres. **d**, Cartoon representation of VRC34-AMC11 SOSIP structure (PDB 6NC3) from same view as c (left) and a different view from above the trimer (right). VRC34 heavy and light chains (PDB 6NC3) are dark and light purple, respectively.

**Table 1.** Data collection and refinement statistics

|  |  |  |
| --- | --- | --- |
|  |  | **Ab1245–BG505–8ANC195** |
| **PDB** |  | **XXXX** |
| **EMD** |  | **xxxx** |
| **Data collection conditions** |  |  |
| Microscope |  | Titan Krios |
| Camera |  | Gatan K3 Summit |
| Magnification |  | 105,000x |
| Voltage (kV) |  | 300 |
| Recording mode |  | counting |
| Dose rate (e<sup>-</sup>/pixel/s) |  | 25 |
| Electron dose (e<sup>-</sup>/Å<sup>2</sup>) |  | 60 |
| Defocus range (µm) |  | 0.8 - 2.5 |
| Pixel size (Å) |  | 0.866 |
| Micrographs collected |  | 2,307 |
| Micrographs used |  | 2,260 |
| Total extracted particles |  | 421,388 |
| Refined particles |  | 172,731 |
| Symmetry imposed |  | C1 |
| Nominal Resolution (Å) |  |  |
|  | FSC 0.5 (unmasked/masked) | 4.0/3.7 |
|  | FSC 0.143 (unmasked/masked) | 3.6/3.5 |
| **Refinement and Validation** |  |  |
| Initial model used |  | 5CJX, 6NC3 |
| Number of atoms |  | 22,171 |
|  | Protein (residues) | 2,634 |
|  | Ligands | BMA:11; NAG:87; MAN:30 |
| MapCC (global/local) |  |  |
| Map sharpening B-factor |  | 171 |
| R.m.s. deviations |  |  |
|  | Bond lengths (Å) | 0.003 |
|  | Bond angles (°) | 0.598 |
| MolProbity score |  | 1.42 |
| Clashscore (all atom) |  | 4.2 |
| Poor rotamers (%) |  | 0.35 |
| Ramachandran plot |  |  |
|  | Favored (%) | 96.59 |
|  | Allowed (%) | 3.37 |
|  | Disallowed (%) | 0.04 |

The Ab1245-BG505-8ANC195 complex structure revealed non-overlapping epitopes at the gp120-gp41 interface for Ab1245 and 8ANC195 (Fig. 2a,b). The Ab1245 Fab was located at the interface between the gp120 and gp41 of its primary Env protomer and the gp41 of a neighboring protomer (Fig. 2c, Fig. 3a). By contrast, 8ANC195 recognized the gp120-gp41 interface of a single protomer with no contacts to neighboring protomers (Fig. 2a), and equivalent interactions with the three BG505 protomers^25^. The Ab1245 epitope overlaps with that of VRC34, a FP-directed bNAb that binds with a three Fab per Env trimer stoichiometry at a site that is located closer to the trimer base^6^ than the Ab1245 epitope (Fig. 2d; Supplementary Fig. 2a). The Ab1245 heavy chain made the majority of contacts with BG505 (only three light chain residues contact BG505), and all but one of the 19 heavy and light chain contact residues were located within CDRs rather than antibody framework regions (FWRs) (Fig. 1a, Fig. 3b). The Ab1245 CDRH1 and CDRH2 loops formed extensive interactions with a portion of gp120 between residues Pro79_gp120_ and Glu87_gp120_ (Fig. 3c), while the long (24-amino acid) CDRH3 loop contacted gp41 residues as well as residues at the termini of gp120 that fit inside the previously-defined membrane-proximal collar^14^ (Fig. 3d). The Ab1245 light chain made contacts with the terminal helix of an adjacent gp41 subunit, which had undergone a change in conformation from a helix to a unstructured region that was partially disordered, suggesting the possibility that the gp41 helix conformation sterically interferes with binding, as was also proposed for the interaction of the human bNAb 3BC315 with BG505 Env trimer^28^ (Fig. 3a,e). The majority of the Ab1245 Fab contacts with BG505 were contacts to protein residues, with the only glycan contact involving the third framework region of the antibody heavy chain (FRWH3) with a terminal sugar on the Asn448_gp120_ glycan (Fig. 3a). By contrast, the 8ANC195 epitope includes required contacts with glycans attached to residues Asn276_gp120_ and Asn234_gp120_^25,29^, and the VRC34 epitope includes contacts with glycans at residues Asn88_gp120_ and Asn241_gp120_^6^ (Fig. 2d). The possibility that an N-glycan attached to Asn611_gp41_ could occlude Ab1245 Fab binding was suggested by a lack of density for this glycan on the primary Ab1245-binding protomer compared with density for one GlcNAc attached to the Asn611_gp41_ residues on the other two protomers (Supplementary Fig. 2b). However, only one Ab1245 Fab bound to a soluble BG505_N611A_ trimer (Fig. 1d), implying that the presence of the Asn611_gp41_ glycan does not account for sub-stoichiometric binding of Ab1245 to Env trimers.

**Figure 3.**
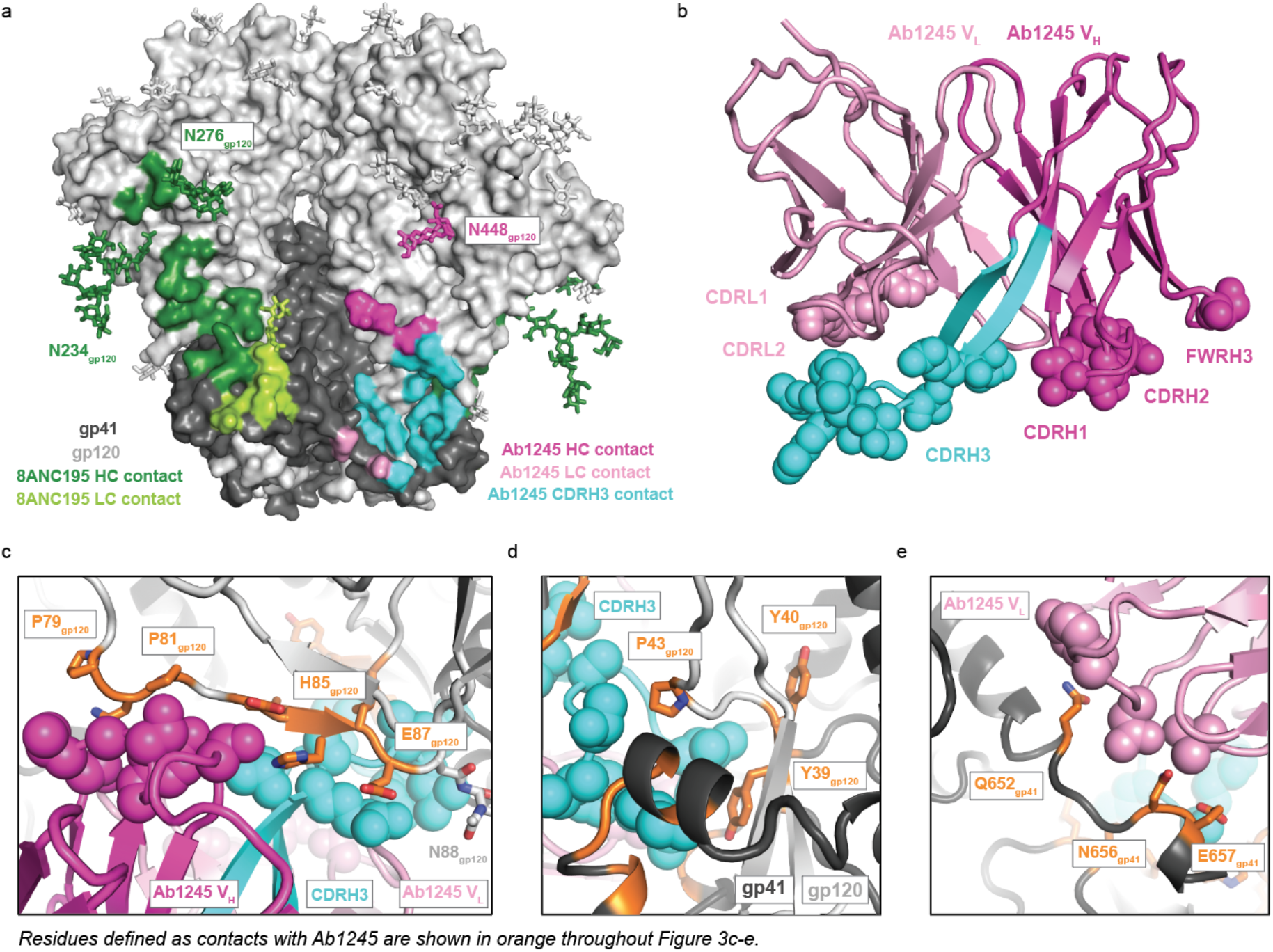
Ab1245 CDRH3 makes the majority of contacts to BG505 Env. **a**, Surface representation of BG505 trimer with colored highlights showing the epitopes of Ab1245 (light and dark pink with CDRH3 contacts highlighted in cyan and pink sticks for glycan contacts) and 8ANC195 (green, with green sticks representing glycans within the epitope). Glycans represented that are not part of an epitope are shown as gray sticks. **b**, Cartoon representation of Ab1245 with paratope residue atoms shown as colored spheres. **c**, gp120 interactions with Ab1245 CDRH1 and CDRH2 loops (dark pink spheres). gp120 is gray with contacts to Ab1245 highlighted in orange. Sidechains of gp120 contacts are shown as sticks. The Ab1245 paratope is represented as in panel b. **d**, Interactions of Ab1245 CDRH3 (cyan spheres) with gp120 (light gray) and gp41 (dark gray). Contacting residues are orange, and side chains discussed in the text are shown. **e**, Ab1245 light chain (light pink spheres) contacts with the terminal helix of an adjacent gp41. Contacting residues are orange with side chains shown.

### The Ab1245 CDRH3 contains a gp41 mimicry motif

Unlike binding of the FP-specific bNAb VRC34^6^ or any other reported HIV-1 antibody, Ab1245 binding to BG505 Env trimer resulted in displacement of the FP (residues 512_gp41_-527_gp41_) and FPPR (residues 528_gp41_-540_gp41_) of the gp41 subunit within the primary protomer to which Ab1245 was bound (Fig. 4a,b). The FP/FPPR displacement resulted from intercalation of the Ab1245 CDRH3 (Fig. 4a). Although the gp41 residues of HR1N (547_gp41_-568_gp41_) are usually disordered in structures of Env trimer (except when an interface antibody is bound), residues N-terminal to this region, 520_gp41_-546_gp41_, are ordered whether or not the Env was complexed with a gp120-gp41 interface antibody (e.g.,^30,31^). In the Ab1245-BG505 complex structure, there was no observed density for residues spanning 512_gp41_-565_gp41_ on the primary gp41 to which Ab1245 was bound, thus both regions of gp41 were disordered. The disorder resulted from Ab1245 binding because residues 520_gp41_ to 546_gp41_ were resolved in the two adjacent gp41 subunits (Fig. 4b).

**Figure 4.**
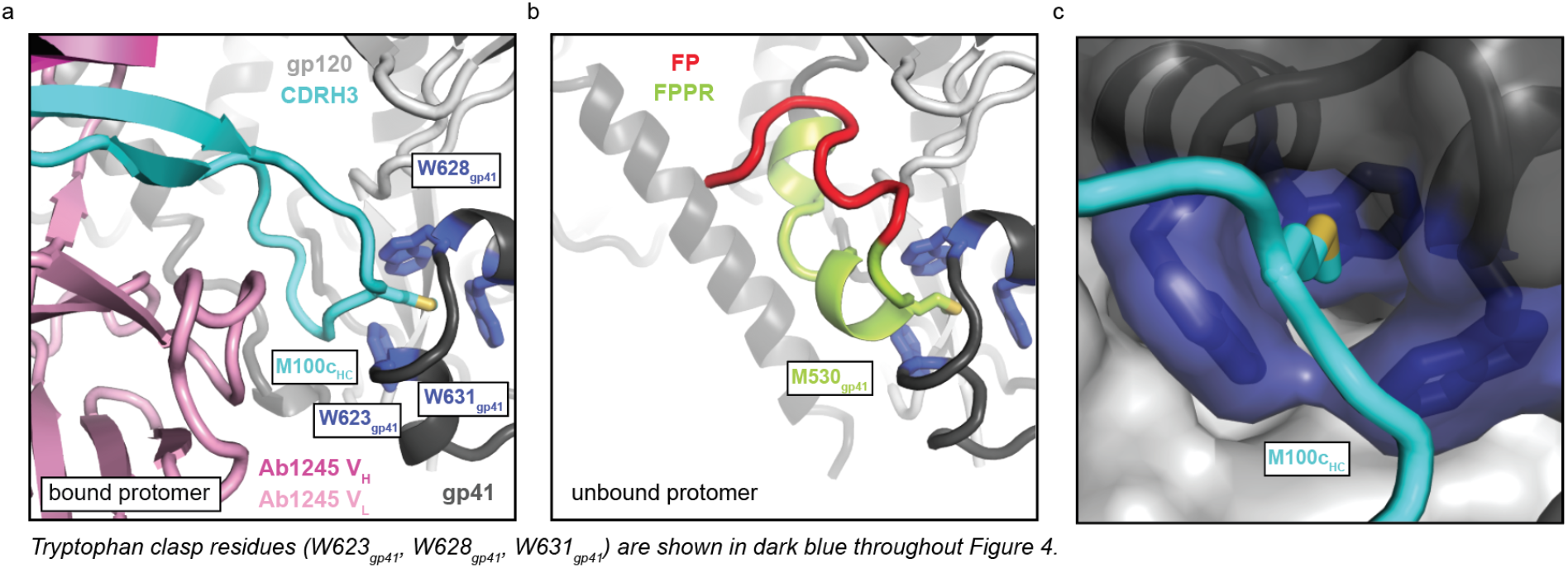
Ab1245 CDRH3 mimics gp41 interactions with the tryptophan clasp. **a**, Cartoon representation of the interactions between Ab1245 V_H_-V_L_ domains (pink with CDRH3 in cyan) and the tryptophan clasp of gp41 (gp41 in dark gray with Trp residues 623_gp41_, 628_gp41_, and 631_gp41_ in dark blue) with gp120 in light gray. Interacting residues between M100c_HC_ and the tryptophan clasp shown as sticks. **b**, Cartoon representation of the same view of an unbound protomer of gp41 with the portion of gp41 containing the fusion peptide (red) and fusion peptide proximal region (FPPR, green) interacting with the gp41 tryptophan clasp (same coloring as in a). Interactions between M530_gp41_ and the tryptophan clasp are shown as sticks. **c**, Surface representation of gp41 (dark gray) and gp120 (light gray) with tryptophan clasp residues in dark blue. Ab1245 CDRH3 is cyan with a stick representation for the Met100c_HC_ sidechain.

One of the disordered gp41 residues in the Ab1245-BG505-8ANC195 structure, Met530_gp41_, normally inserts into the gp41 ‘tryptophan clasp’ formed by residues Trp623_gp41_, Trp628_gp41_, and Trp631_gp41_^14,15^ (Fig. 4a). The tryptophan clasp has been hypothesized to be a key interaction that stabilizes the Env trimer in both its closed, prefusion conformation and its CD4-bound open conformation, and it has been speculated that the disengagement of Met530_gp41_ from the tryptophan clasp triggers elongation of HR1 into a full-length helix and the large rearrangement of the FP required for insertion into the host membrane^14,15^. However, the primary protomer to which Ab1245 is bound contained a disengaged Met530_gp41_, but adopted most structural characteristics of a closed, prefusion Env trimer (Fig. 2a,b). The limited structural changes in Ab1245-bound BG505 from other closed Env trimer structures included disorder of the gp41 FP and FPPR regions, a transition to an unstructured secondary structure in a gp41 terminal helix, a shift in the N88_gp120_ glycan, and absence of density for the N611_gp41_ glycan (Fig. 4a; Supplementary Fig. 2c). The stability of the Ab1245-bound BG505 trimer in the closed conformation despite displacement of Met530_gp41_ from the tryptophan clasp is rationalized by the insertion of an Ab1245 CDRH3 residue, Met100c_1245 HC_, into the gp41 tryptophan clasp to mimic Env residue Met530_gp41_ (Fig. 4a,c).

Some HIV-1 gp120-gp41 interface-binding antibodies induce dissociation of Env trimers into protomers after incubation for 30 minutes to several hours^28,32^. Disruption of residue(s) within the gp41 tryptophan clasp was hypothesized to be the mechanism by which these antibodies induce trimer dissociation^28,33,34^. Indeed, Met530_gp41_, which is part of the disordered gp41 region that is displaced by Ab1245 binding, has been implicated as an important anchoring residue that mediates gp41 dynamics^35^. However, we found no dissociation of Env trimers into protomers in the cryo-EM structure of the Ab1245-BG505-8ANC195 complex that was derived after an overnight incubation of BG505 with Fabs or indication of dissociated protomers by SEC-MALS (Fig. 1c). We hypothesize that disruption of the tryptophan clasp by Ab1245 does not induce trimer dissociation because insertion of its CDRH3 residue Met100c_1245 HC_ mimics gp41 Met530_gp41_ to stabilize the tryptophan clasp conformation (Fig. 4a,b).

### Ab1245 IgG is non-neutralizing

Although Ab1245 Fab exhibits strong binding to BG505 SOSIP Env (Fig. 1b), we observed no neutralization of a BG505 pseudovirus at Ab1245 IgG concentrations up to 1 mg/mL (Fig. 1e). Additionally, we observed no neutralization by Ab1245 Fab at concentrations up to 100 μg/mL for eight strains of pseudovirus (unpublished data).

A recent report described RM20E1, an antibody isolated from BG505-immunized NHPs, which neutralized an N611A_gp41_ mutant strain of BG505, but not the wild-type version of BG505^36^. In common with Ab1245, RM20E1 binds BG505 SOSIP sub-stoichiometrically at an epitope that overlaps with the Ab1245 epitope, but it does not displace the FP or FPPR. To determine whether the Asn611_gp41_ glycan interferes with Ab1245 neutralization of BG505 or viral strains containing a glycan at this position, we also evaluated neutralization of Ab1245 against a BG505_N611A_ strain and eight other HIV-1 strains that lack a PNGS at position 611_gp41_. We observed no neutralization by Ab1245 IgG against any of the viral strains under conditions in which a positive control IgG exhibited neutralization (Fig. 1e).

The fact that the Ab1245 was elicited by a SOSIP immunogen implies that non-neutralizing gp41-mimicking antibodies could be raised during other vaccination experiments. Important features of Ab1245 that allow its gp41 mimicry include a long CDRH3 with a methionine roughly in the middle (Met100c_1245 HC_ is the 11^th^ residue of the 24 amino acid CDRH3). A search of the Protein Data Bank^37^ for Fab structures with features of the Ab1245 CDRH3 (22-26 residue length and a methionine at position 7-13) revealed three of 1657 Fabs with these features (PDBs 5CEZ, 6E4X, and 2XTJ). However, the conformations of the CDRH3s of these Fabs did not resemble the Ab1245 CDRH3 conformation (data not shown). Given that the features required for Ab1245 mimicry of gp41 are apparently rare (i.e., not found in a survey of other antibody structures), Ab1245 CDRH3 characteristics (~24 residue CDRH3 with a methionine at position ~11) could be used to screen elicited antibody sequences to identify Ab1245-like antibodies that are likely to be non-neutralizing. In addition, the Ab1245-BG505 complex structure could inform the engineering of SOSIP immunogens to prevent displacement of the FP and FPPR residues surrounding Met530_gp41_.

Our results suggest that Ab1245 binds to at least some strains of Env trimer (e.g., BG505) on virions, but does not affect entry into target cells. Since only one Ab1245 Fab binds per Env trimer at a site that would not disrupt CD4 binding, Env trimers should still be able to undergo CD4-induced conformational changes^38–41^ allowing coreceptor binding and subsequent insertion of one or two of the trimer FPs into the host cell membrane. This would imply that fusion requires only up to two of three FPs to be inserted into the target membrane. However, it is also possible that the third FP, which was displaced by Ab1245 binding and is disordered in the Ab1245-BG505 structure (Fig. 4a), could access the host cell membrane and insert itself despite Ab1245 binding, thus enabling all three FPs per Ab1245-bound Env trimer to function in membrane fusion.

The characterization of Ab1245 reported here suggest that, in order for an antibody against the FP epitope to be neutralizing, it must directly bind the FP in order to prevent it from inserting into the host cell membrane. Thus displacement of the FP and FPPR by an antibody such as Ab1245 through gp41 mimicry need not result in neutralization. However, by blocking neutralizing antibodies such as VRC34, an antibody that directly interacts with the FP^6^, from binding HIV-1 Envs, Ab1245 and similar antibodies could serve as decoys that protect a conserved epitope on Env from binding neutralizing antibodies.

## Methods

### Single B cell sorting and antibody sequencing

Cells from lymph node biopsies obtained from immunized macaques were thawed and washed in RPMI medium 1640 (1x) (Gibco #11875-093). Macaque cells were incubated with 100 μl of FACS buffer (PBS 1x with 2% fetal bovine serum and 1mM EDTA) with human Fc Block (BD Biosciences #564219) at a 1:500 dilution for 30 min on ice.

BG505 and B41 tetrameric baits were prepared by incubating 5 μg of AviTagged and biotinylated BG505 and B41 SOSIP trimers with fluorophore-conjugated streptavidin at a 1:200 dilution in 1xPBS for 30 min on ice as previously reported^4,42^. Tetramers were mixed with the following anti-human antibody cocktail: anti-CD16 APC-eFluor780 (Invitrogen, #47-0168-41), anti-CD8α APC-eFluor780 (Invitrogen, #47-0086-42), anti-CD3 APC-eFluor780 (Invitrogen, #47-0037-41), anti-CD14 APC-eFluor780 (eBiosciences, #47-0149-41), anti-CD20 PeCy7 (BD, #335793), anti-CD38 FITC (Stem Cell technologies, #60131FI) at a 1:200 dilution and the live/dead marker Zombie NIR at a 1:400 dilution in FACS buffer.

Zombie NIR-/CD16-/CD8a-/CD3-/CD14-/CD20+/CD38+/double BG505^+^ and B41^+^ single cells were isolated from the macaque cell homogenates using a FACS Aria III (Becton Dickinson). Single cells were sorted into individual wells of a 96-well plate containing 5 μl of lysis buffer (TCL buffer (Qiagen #1031576) with 1% of 2-b-mercaptoethanol). Plates were immediately frozen on dry ice and stored at −80°C.

Antibody sequencing and cloning were performed as previously described^4^. Assignments of V, D, and J genes, percent mutated from germline sequences, and identification of CDR loops for Fig. 1a were done using IMGT/V-QUEST analysis using genes from the species macaca mulatta^43–45^. Percent change from germline does not include a one-amino acid insertion in the Ab1245 heavy chain. Residues were numbered according to the Kabat convention^46^.

### Protein Expression

Fabs (Ab1245, 8ANC195, BG1) and IgGs (Ab1245, N6) were expressed and purified as described^47^. Briefly, IgGs and 6xHis-tagged Fabs were expressed by transient transfection of paired heavy chain and light chain expression plasmids into HEK293-6E or Expi293 cells (Life Technologies). Fabs and IgGs were purified from transfected cell supernatants using Ni-NTA (GE Healthcare) (for Fabs) or protein A (GE Healthcare) (for IgG) affinity chromatography followed by SEC on a Superdex 200 16/60 column (GE Healthcare). Proteins were stored in 20 mM Tris, pH 8.0, and 150 mM sodium chloride (TBS buffer).

BG505 SOSIP.664, a soluble clade A gp140 trimer that includes ‘SOS’ substitutions (A501C_gp120_, T605C_gp41_), the ‘IP’ substitution (I559P_gp41_), the N-linked glycan sequence at residue 332_gp120_ (T332N_gp120_), an enhanced gp120-gp41 cleavage site (REKR to RRRRRR), and a stop codon after residue 664_gp41_ (Env numbering according to HX nomenclature)^12^ was expressed in a stable CHO cell line (kind gift of John Moore, Weill Cornell Medical College) as described^48^. BG505_N611A_ SOSIP was expressed by transient transfection in Expi-293 cells as described^25^. SOSIP proteins were isolated from cell supernatants using a 2G12 immunoaffinity column made by covalently coupling 2G12 IgG monomer to an NHS-activated Sepharose column (GE Healthcare). Protein was eluted with 3M MgCl2 followed by buffer exchange into TBS, and trimers were purified using Superdex 200 16/60 SEC (GE Healthcare), and then stored in TBS.

### SEC-MALS

Purified BG5505 SOSIP and BG505-Fab complexes were characterized by SEC-MALS to determine absolute molecular masses^49^. For complexes, BG505 SOSIP.664 was mixed with a 3-fold molar excess of Ab1245 Fab or 8ANC195 Fab relative to BG505 trimer in TBS.

Complexes were incubated overnight at room temperature and injected onto a Superdex 200 10/300 GL gel-filtration chromatography column equilibrated with TBS. The chromatography column was connected with an 18-angle light-scattering detector (DAWN HELEOS II; Wyatt Technology), a dynamic light-scattering detector (DynaPro Nanostar; Wyatt Technology), and a refractive index detector (Optilab t-rEX; Wyatt Technology). Data were collected every second at 25°C at a flow rate of 0.5 mL/min. Calculated molecular masses were obtained by data analysis using the program ASTRA 6 (Wyatt Techonology).

### Mass photometry

Microscope coverslips (No. 1.5, 24 × 50 mm, VWR) were cleaned by sequential rinsing with Milli-Q H2O followed by isopropanol and again Milli-Q H2O followed by drying using a filtered pressured air stream. Clean coverslips were assembled using CultureWell™ reusable silicon gaskets (Grace Bio-Labs, # 103250). Measurements were performed using a OneMP mass photometer (Refeyn Ltd, Oxford, UK). Immediately before each measurement, wells were filled with 15 μl TBS buffer. The focal position was identified and secured in place with an autofocus system based on total internal reflection for the entire measurement. Immediately following the focusing procedure, 1 μl of protein solution was added and gently mixed by pipetting up and down 3 times at a 5 μl mixing volume. Calibration standards (1 μM Bovine Serum Albumin (BSA) SIGMA #23209 and 500 nM apoferritin SIGMA #A3360) were measured first. SOSIP-Fab complexes (incubated at 7 μM SOSIP and ~7.7 μM Fab for either 2-5 days or for 1-2 hours and subsequently diluted 1:6 in TBS) was added to the 15 μl PBS buffer in the well resulting in an ~114 nM concentration with respect to the SOSIP unless indicated otherwise. Recording of a mass photometry movie was started immediately. Data acquisition was performed using AcquireMP 2.2.0 software (Refeyn Ltd.), and data analysis was carried out using DiscoverMP 2.2.0 software (Refeyn Ltd.). Resulting mass photometry graphs were evaluated and protein complex masses were determined against the BSA/apoferritin calibration measurements carried out on the same day.

### Complex formation and cryo-EM data collection

Ab1245-BG505 complex was prepared by adding a 3-fold molar excess of Ab1245 Fab (Fab to BG505 protomer) to CHO-expressed BG505 SOSIP.664 isolated from the second half of a monodisperse SEC peak. This mixture was incubated at room temperature for three hours, after which a 3-fold molar excess of 8ANC195 Fab to BG505 protomer was added and the complex was incubated at room temperature overnight. The Ab1245-BG505-8ANC195 complex was then purified using size-exclusion chromatography (Superdex 200) and concentrated to 4.7 mg/mL before vitrification on a freshly-glow discharged (15 mA for 1 min, Ted Pella PELCO easiGLOW) Quantifoil R 2/2 300 mesh Cu grid (Electron Microscopy Services). Samples were vitrified using a Mark IV Vitrobot (Thermo Fisher) in 100% liquid ethane after a 3 second blot with Whatman No. 1 filter paper at 22°C and 100% humidity. Micrograph movies were collected on a Titan Krios using SerialEM^50^ automated collection software with a K3 camera (Gatan) operating in super resolution mode at a nominal magnification of 105,000x (0.433 Å/ pixel) using a defocus range of −0.8 to −2.5 μm. The dose was 1.5 e-/Å^2^ over 40 frames, resulting in a total dose of 60 e-/Å^2^. Data collection conditions are summarized in Supplementary Table 1.

### Cryo-EM data processing

Processing was carried out entirely within Relion-3^51,52^. First, micrograph movies were motion corrected, dose-weighted, and binned to 0.866 Å/pixel using Motioncor2^53^, and then the nondose-weighted micrographs were used for CTF estimation using Gctf^54^. Micrographs with poor CTF fits or signs of crystalline ice were discarded. Selected micrographs then underwent autopicking after which 4×4 binned particles were extracted (3.46 Å/pixel). These particles were then subjected to reference-free 2D classification after which selected particles underwent three rounds of iterative 3D classification, wherein classes representing 8ANC195-BG505 were discarded and only the final particles representing 1245-BG505-8ANC195 were selected and unbinned (0.866 Å/pixel). Finally, these unbinned particles underwent 3D refinement (C1 symmetry imposed) and were post-processed into a map with a gold-standard FSC calculation^55^ of 3.7 Å. A ‘blurred’ map was also created using a higher B-factor to uncover N-linked glycan densities.

### Model building

Coordinates of BG505-8ANC195 Fab V_H_-V_L_ domains (PDB 5CJX) and VRC34 Fab V_H_-V_L_ domains (PDB 6NC3) were fitted into map density using UCSF Chimera^56^. Coordinates were then built into densities using iterative rounds of refinement in Phenix^57^ (rigid body and real-space refinement) and Coot^58^. Antibody numbering was done in the Kabat convention using the online ANARCI server^59^.

### Structural analysis

Structure figures were made using UCSF Chimera^56^ or PyMol^60^. Contact residues were assigned as residues with any atom located <4.0 Å from an atom in a residue on the partner molecule. Hydrogen bond interactions were not assigned due to limited resolution.

### In vitro neutralization assays

Pseudovirus neutralization assays were conducted as described^61^ either in-house (for strains BG505, BG505_N611A_, CE0217, CNE55, JRCSF, Du422, T250-4, Tro, X1632, 246F3, CH119, CE1176, BJOX002000_03_02, 25710, X2278, CNE8, and 398F1) or by the Collaboration for AIDS Vaccine Discovery (CAVD) core neutralization facility (for the remaining strains in Fig. 1e). IgGs (Ab1245 and an N6^62^ positive control) were evaluated in duplicate with an 8-point, 4-fold dilution series starting at a top concentration of 500 or 1000 μg/mL for in-house neutralizations or an 8-point, 5-fold dilution series starting at a top concentration of 250 μg/mL at the CAVD facility.

## Data Availability

The atomic model and and cryo-EM maps have been deposited in the Protein Data Bank (PDB) accession code XXXX and Electrion Microscopy Data Bank (EMDB) entry EMD-XXXXX.

## Acknowledgements

We thank Anthony P. West for help with analysis of antibody CDRH3 sequences. Cryo-EM was performed in the Beckman Institute Resource Center for Transmission Electron Microscopy at Caltech with assistance from directors A. Malyutin and S. Chen. We thank the Beckman Institute Protein Expression Center at Caltech for protein production, John Moore (Weill Cornell Medical College) for the BG505 stable cell line, Gabriella Kiss, Sofia Ferreira, and Brenda Watt at Refeyn Ltd. for providing a demonstration model OneMP mass photometer, training, and materials to Caltech, Kristie M. Gordon (The Rockefeller University) for assistance with flow cytometry, and Rogier W. Sanders and Marit J. van Gils (Academisch Medisch Centrum Universiteit van Amsterdam) for providing AviTagged and biotinylated BG505 and B41 SOSIP trimers. This work was supported by the National Institute of Allergy and Infectious Diseases (NIAID) HIVRAD P01 AI100148 (to P.J.B. and M.C.N.), Gates CAVD grant INV-002143 (to P.J.B., M.C.N., and M.A.M.), an NSF Graduate Research Fellowship (to M.E.A.), and a Bill and Melinda Gates Foundation grant (#OPP1146996 to M.S.S.). M.C.N. is an HHMI Investigator.

## Author contributions

M.E.A. and P.J.B. designed the research. M.E.A., H.B.G., J.V., J.R.K., and Y.E.L. performed biophysical experiments. M.E.A., H.B.G., J.V., J.R.K., and P.J.B. analyzed the results. P.NP.G. and M.S.S. carried out and supervised *in vitro* neutralization assays. A.E. and M.C.N. carried out and supervised the derivation of monoclonal antibody sequences and plasmids from NHPs. R.G. and M.A.M. planned and supervised the immunization experiments in NHPs. M.E.A and P.J.B. wrote the manuscript with input from co-authors.

---

**Supplementary Fig. 1.**
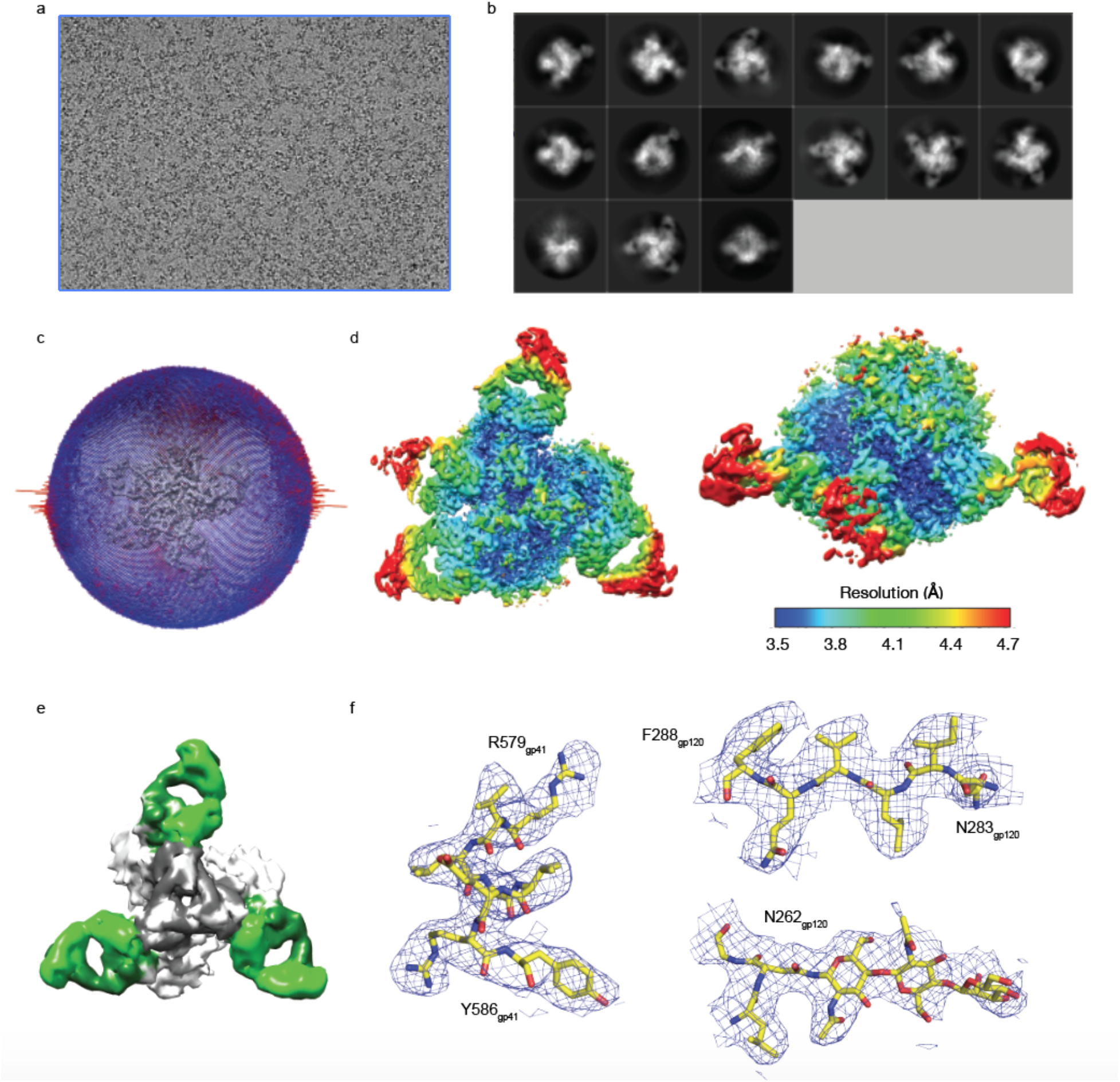
Cryo-EM Data Processing. **a**, Representative motion-corrected micrograph. **b**, 4x-binned selected 2D classes. **c**, Orientation distribution analysis. **d**, Local resolution map from two angles. **e**, Map of 3D class that contained 3 8ANC195 Fabs and no Ab1245 Fabs (colors as in Fig. 2b). **f**, Representative images of map quality at indicated sites for the 3.7 Å Ab1245-BG505-8ANc195 cryo-EM structure.

**Supplementary Fig. 2.**
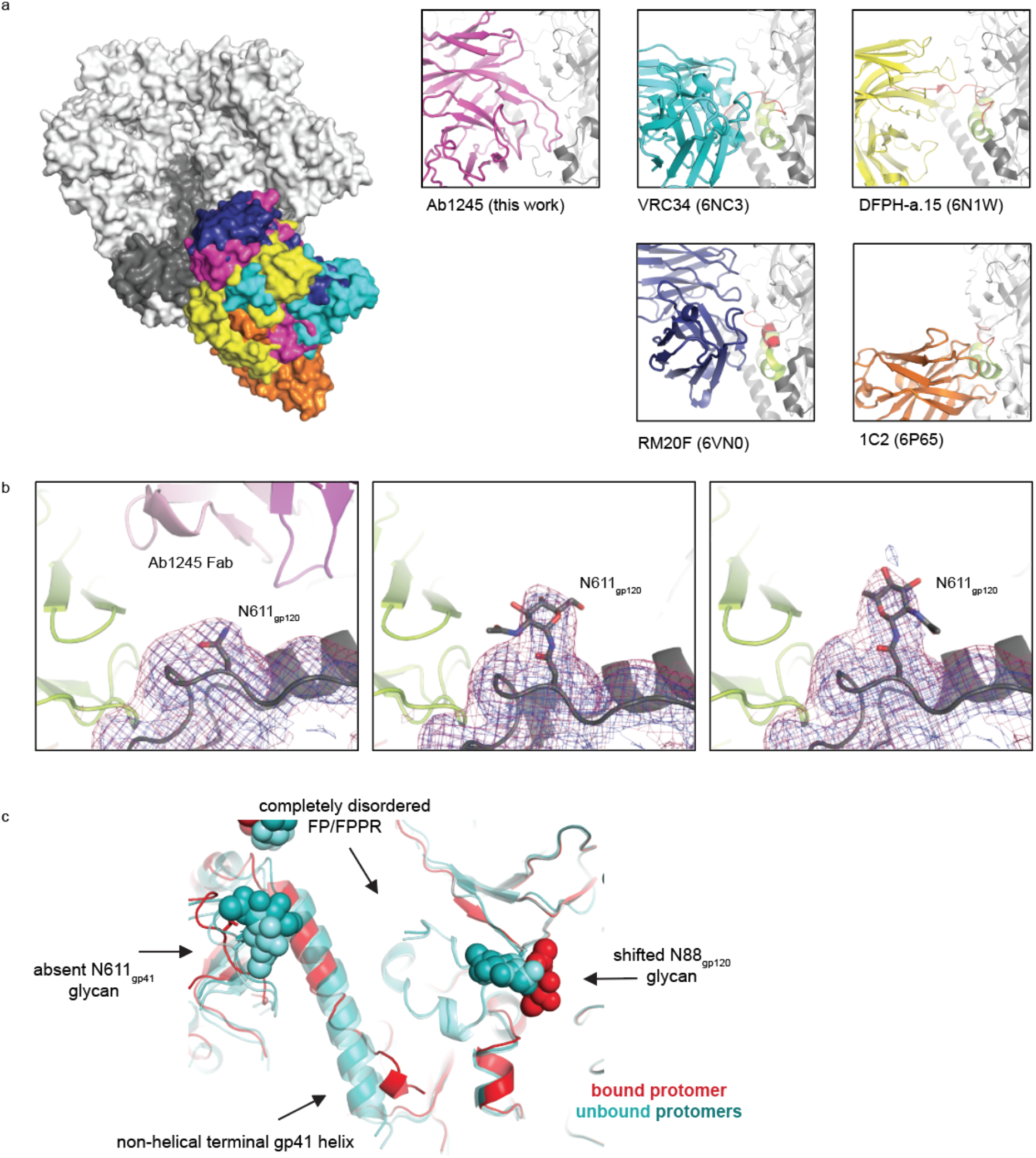
Ab1245 Structural Analysis. **a**, Comparison of binding of gp120-gp41 interface or fusion peptide-directed antibodies. Left: BG505 trimer (gray surface) and Fab V_H_-V_L_ domains (colored surfaces). Insets: individual antibodies bound to Env trimer (gray) are shown as cartoon with the FP in red and the FPPR in orange. **b**, Comparison of density for the N611_gp41_ glycan on gp120 (gray cartoon), with 8ANC195 (green cartoon), and Ab1245 (pink cartoon with non-blurred (blue) and blurred (red) maps shown as mesh). **c**, Comparison of secondary structure between protomers of gp120-gp41 with Ab1245 bound (red cartoon) and not bound (shades of teal, cartoon). Glycans are shown as spheres, and differences between bound and unbound protomers are highlighted with arrows and descriptions.

## Uncategorized References

1. Andrabi, R., Bhiman, J.N. & Burton, D.R. Strategies for a multi-stage neutralizing antibody-based HIV vaccine. Curr Opin Immunol 53, 143–151 (2018).

2. Escolano, A., Dosenovic, P. & Nussenzweig, M.C. Progress toward active or passive HIV-1 vaccination. J Exp Med 214, 3–16 (2017).

3. Kwong, P.D. & Mascola, J.R. HIV-1 Vaccines Based on Antibody Identification, B Cell Ontogeny, and Epitope Structure. Immunity 48, 855–871 (2018).

4. Escolano, A. et al. Immunization expands B cells specific to HIV-1 V3 glycan in mice and macaques. Nature 570, 468–473 (2019).

5. Steichen, J.M. et al. A generalized HIV vaccine design strategy for priming of broadly neutralizing antibody responses. Science 366 (2019).

6. Kong, R. et al. Fusion peptide of HIV-1 as a site of vulnerability to neutralizing antibody. Science 352, 828–33 (2016).

7. Xu, K. et al. Epitope-based vaccine design yields fusion peptide-directed antibodies that neutralize diverse strains of HIV-1. Nat Med 24, 857–867 (2018).

8. Jardine, J. et al. Rational HIV immunogen design to target specific germline B cell receptors. Science 340, 711–6 (2013).

9. McGuire, A.T. et al. Specifically designed immunogens select and activate B cells expressing precursors of broadly neutralizing human antibodies to HIV-1 in knock-in mice. Nature Comm 7, 10618 (2016).

10. Bianchi, M. et al. Electron-Microscopy-Based Epitope Mapping Defines Specificities of Polyclonal Antibodies Elicited during HIV-1 BG505 Envelope Trimer Immunization. Immunity 49, 288–300 e8 (2018).

11. McCoy, L.E. et al. Holes in the Glycan Shield of the Native HIV Envelope Are a Target of Trimer-Elicited Neutralizing Antibodies. Cell Rep 16, 2327–38 (2016).

12. Sanders, R.W. et al. A next-generation cleaved, soluble HIV-1 Env Trimer, BG505 SOSIP.664 gp140, expresses multiple epitopes for broadly neutralizing but not non-neutralizing antibodies. PLoS Pathog 9, e1003618 (2013).

13. Wang, Z. et al. Isolation of single HIV-1 Envelope specific B cells and antibody cloning from immunized rhesus macaques. J Immunol Methods 478, 112734 (2020).

14. Pancera, M. et al. Structure and immune recognition of trimeric pre-fusion HIV-1 Env. Nature 514, 455–61 (2014).

15. Burton, D.R. & Hangartner, L. Broadly Neutralizing Antibodies to HIV and Their Role in Vaccine Design. Annu Rev Immunol 34, 635–59 (2016).

16. Steichen, J.M. et al. HIV Vaccine Design to Target Germline Precursors of Glycan-Dependent Broadly Neutralizing Antibodies. Immunity 45, 483–96 (2016).

17. Duan, H. et al. Glycan Masking Focuses Immune Responses to the HIV-1 CD4-Binding Site and Enhances Elicitation of VRC01-Class Precursor Antibodies. Immunity 49, 301–311 e5 (2018).

18. Garrity, R.R. et al. Refocusing neutralizing antibody response by targeted dampening of an immunodominant epitope. J Immunol 159, 279–89 (1997).

19. Klasse, P.J. et al. Epitopes for neutralizing antibodies induced by HIV-1 envelope glycoprotein BG505 SOSIP trimers in rabbits and macaques. PLoS Pathog 14, e1006913 (2018).

20. Brune, K.D. et al. Plug-and-Display: decoration of Virus-Like Particles via isopeptide bonds for modular immunization. Sci Rep 6, 19234 (2016).

21. Zakeri, B. et al. Peptide tag forming a rapid covalent bond to a protein, through engineering a bacterial adhesin. Proc Natl Acad Sci U S A 109, E690–7 (2012).

22. Vigdorovich, V. et al. Repertoire comparison of the B-cell receptor-encoding loci in humans and rhesus macaques by next-generation sequencing. Clin Transl Immunology 5, e93 (2016).

23. Wang, H. et al. Asymmetric recognition of HIV-1 Envelope trimer by V1V2 loop-targeting antibodies. Elife 6 (2017).

24. Scharf, L. et al. Antibody 8ANC195 reveals a site of broad vulnerability on the HIV-1 envelope spike. Cell Rep 7, 785–95 (2014).

25. Scharf, L. et al. Broadly Neutralizing Antibody 8ANC195 Recognizes Closed and Open States of HIV-1 Env. Cell 162, 1379–90 (2015).

26. Sonn-Segev, A. et al. Quantifying the heterogeneity of macromolecular machines by mass photometry. Nat Commun 11, 1772 (2020).

27. Scheid, J.F. et al. Sequence and Structural Convergence of Broad and Potent HIV Antibodies That Mimic CD4 Binding. Science 333, 1633–1637 (2011).

28. Lee, J.H. et al. Antibodies to a conformational epitope on gp41 neutralize HIV-1 by destabilizing the Env spike. Nat Commun 6, 8167 (2015).

29. West, A.P., Jr. et al. Computational analysis of anti-HIV-1 antibody neutralization panel data to identify potential functional epitope residues. Proc Natl Acad Sci U S A 110, 10598–603 (2013).

30. Gristick, H.B. et al. Natively glycosylated HIV-1 Env structure reveals new mode for antibody recognition of the CD4-binding site. Nat Struct Mol Biol 23, 906–915 (2016).

31. Schommers, P. et al. Restriction of HIV-1 Escape by a Highly Broad and Potent Neutralizing Antibody. Cell 180, 471–489 e22 (2020).

32. Dubrovskaya, V. et al. Targeted N-glycan deletion at the receptor-binding site retains HIV Env NFL trimer integrity and accelerates the elicited antibody response. PLoS Pathog 13, e1006614 (2017).

33. Dubrovskaya, V. et al. Vaccination with Glycan-Modified HIV NFL Envelope Trimer-Liposomes Elicits Broadly Neutralizing Antibodies to Multiple Sites of Vulnerability. Immunity 51, 915–929 e7 (2019).

34. Turner, H.L. et al. Disassembly of HIV envelope glycoprotein trimer immunogens is driven by antibodies elicited via immunization. bioRxiv 10.1101/2021.02.16.431310(2021).

35. Kumar, S. et al. Capturing the inherent structural dynamics of the HIV-1 envelope glycoprotein fusion peptide. Nat Commun 10, 763 (2019).

36. Desrosiers, R.C. et al. Mapping the immunogenic landscape of near-native HIV-1 envelope trimers in non-human primates. PLOS Pathogens 16, e1008753 (2020).

37. Berman, H.M. et al. The Protein Data Bank. Nucleic Acids Research 28, 235–242 (2000).

38. Wang, H. et al. Cryo-EM structure of a CD4-bound open HIV-1 envelope trimer reveals structural rearrangements of the gp120 V1V2 loop. Proc Natl Acad Sci U S A 113, E7151–E7158 (2016).

39. Ozorowski, G. et al. Open and closed structures reveal allostery and pliability in the HIV-1 envelope spike. Nature 547, 360–363 (2017).

40. Wang, H., Barnes, C.O., Yang, Z., Nussenzweig, M.C. & Bjorkman, P.J. Partially Open HIV-1 Envelope Structures Exhibit Conformational Changes Relevant for Coreceptor Binding and Fusion. Cell Host Microbe 24, 579–592 e4 (2018).

41. Yang, Z., Wang, H., Liu, A.Z., Gristick, H.B. & Bjorkman, P.J. Asymmetric opening of HIV-1 Env bound to CD4 and a coreceptor-mimicking antibody. Nat Struct Mol Biol 26, 1167–1175 (2019).

42. Wang, Z. et al. A broadly neutralizing macaque monoclonal antibody against the HIV-1 V3-Glycan patch. eLife 9(2020).

43. Giudicelli, V. & Lefranc, M.P. IMGT/junctionanalysis: IMGT standardized analysis of the V-J and V-D-J junctions of the rearranged immunoglobulins (IG) and T cell receptors (TR). Cold Spring Harb Protoc 2011, 716–25 (2011).

44. Giudicelli, V., Brochet, X. & Lefranc, M.P. IMGT/V-QUEST: IMGT standardized analysis of the immunoglobulin (IG) and T cell receptor (TR) nucleotide sequences. Cold Spring Harb Protoc 2011, 695–715 (2011).

45. Brochet, X., Lefranc, M.P. & Giudicelli, V. IMGT/V-QUEST: the highly customized and integrated system for IG and TR standardized V-J and V-D-J sequence analysis. Nucleic Acids Res 36, W503–8 (2008).

46. Kabat, E.A., Wu, T.T., Perry, H.M., Gottesman, K.S. & Foeller, C. Sequences of proteins of immunolgical interest. Department of Health and Human Services, Washington, D.C. (1991).

47. Scharf, L. et al. Structural basis for germline antibody recognition of HIV-1 immunogens. Elife 5 (2016).

48. Dey, A.K. et al. cGMP production and analysis of BG505 SOSIP.664, an extensively glycosylated, trimeric HIV-1 envelope glycoprotein vaccine candidate. Biotechnol Bioeng 10.1002/bit.26498(2017).

49. Wyatt, P.J. Light scattering and the absolute characterization of macromolecules. Analytica Chimica Acta 272, 1–40 (1993).

50. Mastronarde, D.N. Automated electron microscope tomography using robust prediction of specimen movements. J Struct Biol 152, 36–51 (2005).

51. Scheres, S.H. RELION: implementation of a Bayesian approach to cryo-EM structure determination. J Struct Biol 180, 519–30 (2012).

52. Zivanov, J. et al. New tools for automated high-resolution cryo-EM structure determination in RELION-3. Elife 7(2018).

53. Zheng, S.Q. et al. MotionCor2: anisotropic correction of beam-induced motion for improved cryo-electron microscopy. Nat Methods 14, 331–332 (2017).

54. Zhang, K. Gctf: Real-time CTF determination and correction. J Struct Biol 193, 1–12 (2016).

55. Scheres, S.H. & Chen, S. Prevention of overfitting in cryo-EM structure determination. Nat Methods 9, 853–4 (2012).

56. Goddard, T.D., Huang, C.C. & Ferrin, T.E. Visualizing density maps with UCSF Chimera. J Struct Biol 157, 281–7 (2007).

57. Liebschner, D. et al. Macromolecular structure determination using X-rays, neutrons and electrons: recent developments in Phenix. Acta Crystallogr D Struct Biol 75, 861–877 (2019).

58. Emsley, P., Lohkamp, B., Scott, W.G. & Cowtan, K. Features and development of Coot. Acta Crystallogr D Biol Crystallogr 66, 486–501 (2010).

59. Dunbar, J. & Deane, C.M. ANARCI: antigen receptor numbering and receptor classification. Bioinformatics 32, 298–300 (2015).

60. Schrödinger, L. The PyMOL Molecular Graphics System. 1.2r3pre edn (The PyMOL Molecular Graphics System, 2011).

61. Montefiori, D.C. Evaluating neutralizing antibodies against HIV, SIV, and SHIV in luciferase reporter gene assays. Curr Protoc Immunol Chapter 12, Unit 12 11 (2005).

62. Huang, J. et al. Identification of a CD4-Binding-Site Antibody to HIV that Evolved Near-Pan Neutralization Breadth. Immunity 45, 1108–1121 (2016).

